# Gut lymph purification regulates monocyte activity in rats with ischemia-reperfusion injury-induced sepsis

**DOI:** 10.1101/2020.02.02.930610

**Authors:** Wei Zhang, Jie Chen, Can Jin, Shuncheng Zhang, Juan Gu, Meimei Shi

## Abstract

**Objective:** To confirm that gut lymph purification (GLP) based on oXiris regulates monocyte activity by targeting the removal of ischemia-reperfusion injury (IRI)-induced intestinal toxic substances (ITSs) in rats.

**Methods:** Sepsis was induced by intestinal IRI in 24 adult male Sprague-Dawley rats that were randomly divided into the control, intestinal IRI, and IRI+GLP groups. The gut lymph fluid (GLF) was drained for 180 minutes. The ITSs levels and the proliferation, apoptosis and positive expression rates of MHC-II molecules of monocytes coincubated with the GLF were detected.

**Results:** Endotoxin, TNF-α, IL-4, IL-6 and IL-10 levels in the lymph and plasma of the IRI group were significantly higher than those of the control group (*p* < 0.01). Compared with the IRI group, GLP treatment significantly decreased the ITS levels (*p* < 0.05). Monocyte proliferation and the positive expression rate of MHC-□ molecules were significantly reduced after co-culturing with GLF upon IRI (*p* < 0.01), and the apoptotic rate was significantly increased (*p* < 0.01). However, culturing monocytes with GLP significantly enhanced the monocyte proliferation, increased the positive expression rate of MHC-□ monocytes (*p* < 0.01), and reduced the apoptotic rate (*p* < 0.01).

**Conclusions:** GLP therapy based on oXiris effectively removed ITSs from the GLF after IRI, thereby blocking the main process of multiple organ dysfunction syndrome by regulating monocyte activity.

## Introduction

During the pathophysiology of a critical illness, blood is redistributed to reduce circulation in the gastrointestinal tract as a protective reflex in the acute response, thus resulting in intestinal ischemia-reperfusion injury (IRI). Therefore, the gut is one of the first organs to be injured. Deitch et al. first proposed the gut lymph (GL) theory in 2006.[1] This theory states that pathogen injury to the gut results in IRI and leads to destruction of the intestinal mucosal barrier. Intestinal toxic substances (ITSs) that are first absorbed into mesenteric lymphatic vessels are carried into the subclavian vein through the thoracic duct[2] and finally disseminated into the blood circulation, which activates the monocyte-macrophage system. This successive iteration produces a closed-loop system that contributes to cascade amplification of inflammation. Early studies suggested that the increased intestinal permeability could lead to bacterial translocation and multiple organ dysfunction syndrome (MODS).[3] The latest experimental evidence shows that GL, as a carrier of danger-associated molecular patterns, is transported to the lungs and systemic circulation, leading to lung injury and MODS.[4] Research on traumatic/hemorrhagic shock (T/HS), severe abdominal infections, burns, severe pancreatitis and heatstroke replicated these injuries in animal models of acute lung injury (ALI), acute respiratory distress syndrome, acute kidney injury, and acute myocardial injury by transfusing gut lymph fluid (GLF) from an IRI rat model into the blood of healthy rats.[5–10] Sufficiently draining the GLF or ligating the gut lymph vessels (GLVs or thoracic ducts) significantly reduced the risk of generator organ injury in rats.[11–15] These studies suggest that the GLF plays a pivotal role in MODS pathogenesis. Therefore, the gut is thought to be the origin of inflammation and the mediator of MODS.[16, 17] Theoretically, GL interventions may interrupt the process of initiating IRI in MODS. Among these interventions, both GLF drainage and GLV ligation can achieve this goal. However, these two “anti-physiological” therapies are unfavorable for application in clinical practice. Therefore, the theory of gut lymph purification (GLP) was derived by expanding on the process of blood purification therapy. GL theory states that the GLP is a therapeutic technology in which the GLF is drained out of the animal or human body via thoracic duct puncture and catheterization, then purified using a lymph purification device (LPD) and transfused back into the body. In the LPD, the filtration membrane technology is pivotal and mainly depends on the ITSs in the GLF. OXiris, an adsorbed hemofiltration membrane, is a substitute product based on the hydrogel structure of a propylene and sodium mesylate polymer (AN69), which has been widely applied in recent years to treat sepsis and septic shock.[18, 19] However, its clinical efficacy in GLP remains unclear. This study was conducted to confirm that GLP therapy based on oXiris facilitates monocyte function by targeting ITSs in the GLF of IRI-induced rats.

## Materials and methods

This was an animal validation study to test the clinical application of GLF purification technology. This study conformed to the Animals in Research: Reporting In Vivo Experiments (ARRIVE) guidelines.

### Animals

Healthy adult male Sprague-Dawley rats (specific-pathogen-free grade, weighing 500±50 g) were purchased from Changsha Tianqin Biotechnology Co., Ltd. (Tianjin City, China), with the license number SCXK (Hunan) 2014-0011. Rats were maintained at 22±2□ on a 12-h light/dark cycle. All experiments involving animals were performed in accordance with the guidance of the Animal Care and Use of Laboratory Animals. The Ethics Committee of Experimental Animals and Use of Laboratory Animals of Zunyi Medical University approved all animal experiments. Experimental animals were killed by cervical dislocation at the end of the experiment.

### Adsorption column based on oXiris biofilm technology

The fabrication details of the adsorption column were as follows: The coats were sawed neatly along a 0.25-mL scale using two 1-mL injectors with the core rod removed. The rubber chips adhering to the coats were washed with saline solution and dried. After combining the broken ends of both coats, the oXiris filter wires were filled in. The joint of the coats of the two syringes were pasted across using a pre-cut 1.5-cm-wide one-time medical transparent dressing for compactness. The oXiris adsorption column was sterilized with ethylene oxide, sealed and stored at room temperature.

### Experimental grouping

Twenty-four adult male Sprague-Dawley rats were randomly divided into the control, intestinal IRI and IRI+GLP groups (n=8 per group). Rats in the control group underwent intestinal lymphatic stem puncture and catheterization without clamping the superior mesenteric artery. For the IRI and IRI+GLP groups, the rats underwent intestinal lymphatic stem puncture and catheterization, followed by clamping with a noninvasive vascular clamp for 60 minutes, then loosened for another 120 minutes.

### Jugular venipuncture catheterization

After establishing the GLF drainage model, the rats’ necks were shaved and disinfected with a cotton ball soaked in alcohol. The skin from the right sternoclavicular connection from the midpoint of the neck was incised vertically to 1.5 cm to expose the subcutaneous connective tissue. After gradual blunt separation, the jugular vein was visualized and isolated. After completing this process, we induced congestion and swelling by clamping the proximal and distal vein with a noninvasive vascular clamp, then cut a small aperture in the middle of this section, which was slightly smaller than the external diameter of the silica gel catheter that was preflushed with heparin saline. Finally, within the vein orifice, the vascular clamp was loosened, the silica gel catheter was sutured and fixed, and the connective tissue and skin were sutured.

### GL and plasma collection

The GLF was drained from all rats for 180 minutes. Approximately 600 μL of GLF per sample were collected from the control and IRI groups and centrifuged for 10 minutes at 10,000 ×g at 4□. The supernatant was stored at −80□ for further examination (the latter sample followed the same steps without further elaboration). The remaining lymph was slowly transfused through the jugular vein; the lymph from the IRI+GLP group was purified through the oXiris adsorption column, and the remaining lymph was slowly transfused through the jugular vein. After 30 minutes of lymph reinfusion, 2 mL of venous blood was collected via inferior vena cava puncture into tubes containing a heparin sodium and centrifuged at 1,000 ×g at 4□ for 10 minutes.

### Detection of ITSs in the lymph and plasma

The ITSs, including endotoxin, TNF-α, IL-4, IL-6 and IL-10, in the lymph fluid and plasma were detected via enzyme-linked immunosorbent assay (ELISA) per the manufacturer’s instructions.

### Detection of monocyte proliferation

After isolating the monocytes, the number of monocytes was adjusted to 5×10^4^/mL. The dosing hole, blank hole and 0-dosing hole were pre-arranged in a 96-well plate, which included monocytes, CCK-8 solution and lymph in the drug-adding pore. The blank pore contained complete medium and CCK-8 solution without monocytes. Monocytes and CCK-8 solution with no added gut lymph were added to the corresponding hole containing 100 μL of the monocyte suspension in the 96-well plate. The culture plate was incubated in a 5% CO_2_ incubator at 37□ for 24 hours. Ten microliters of the lymph solutions were added into the corresponding holes of the culture plate and incubated for 20 hours in a 5% CO_2_ incubator at 37□. Next, 10 μL of the CCK-8 solutions were added to each hole without forming bubbles, then the culture plate was incubated in an incubator for 3 hours. The OD value at 450 nm was determined via enzyme labeling. Monocyte proliferation was calculated using the formula (cell viability [%] = [A (medication) - A (blank)]/[A (0 medication) - A (blank)] ×100).

### Detection of apoptosis in monocytes

After isolating the monocytes, the number of monocytes was adjusted to 5×10^4^/mL, and the lymph and cell suspension were mixed at a 1:10 ratio. Next, 1.5 mL of the monocyte suspension was added to each hole of the 6-well plate, then 150 μL of lymph was added to each hole and cultured in a 5% CO_2_ incubator at 37□ for 24 hours. Cells were gently scraped and suspended with 5 mL of phosphate-buffered saline (PBS), then centrifuged at 250 g for 10 minutes. The precipitate was resuspended with 5 mL of PBS and centrifuged for another 10 minutes. Finally, 5 μL of FITC/Annexin V and 5 μL of propidium iodide were added to 4 mL of the suspended cells. Flow cytometry was performed within 1 hour.

### Detection of the monocyte antigen-presenting function

After isolating the monocytes, the number of monocytes was adjusted to 5×10^4^/mL, and 1.5 mL of monocyte suspension was added to each hole in the 6-well plate. Next, 150 μL of lymph was added to each well and cultured in a 5% CO_2_ incubator at 37□ for 24 hours. Cells were scraped and suspended with 5 mL of PBS, then centrifuged at 250 g for 10 minutes. The precipitate was resuspended with 5 mL of PBS and centrifuged for another 10 minutes, then the cells in PBS were resuspended, and 1 μL of OX-6 and 2 μL of OX-42 were added to avoid light for up-flow detection within 1 hour.

Data are presented as the mean ± standard error of the mean (SEM) or the mean ± SEM% and analyzed (not in a blinded manner) using software SPSS 22.0 for Windows. Normality was tested with the Shapiro-Wilk test, and equal variance was evaluated before the analysis of variance (ANOVA). Data were statistically analyzed using Student’s t-test (two-tailed), the Kolmogorov-Smirnov test, one-way ANOVA, or repeated-measures ANOVA, as stated individually in the results section. The ANOVAs were followed by Fisher’s post hoc tests. Statistical significance was set at *p* < 0.05.

## Results

### ITS levels in the blood and lymph

ITS levels (endotoxin, TNF-a, IL-4, IL-6 and IL-10) in the lymph and plasma of all rats were detected via ELISA, and the levels were significantly increased after IRI treatment (*p* < 0.01). Compared with the IRI group, GLP therapy significantly decreased the levels of endotoxin, TNF-α, IL-4, IL-6 and IL-10 in the lymph fluid (*p* < 0.05) but did not affect the levels of endotoxin, TNF-α, IL-4, IL-6 and IL-10 in the blood (*p* > 0.05; Tables 1, 2 and 3).

### Monocytes

Monocytes were cultured for 2 to 4 hours. The monocytes were round, oval, kidney-shaped, horseshoe-shaped, or irregular (Fig 2). Trypan blue staining showed that more than 95% of the cells were living.

**Fig 1.**
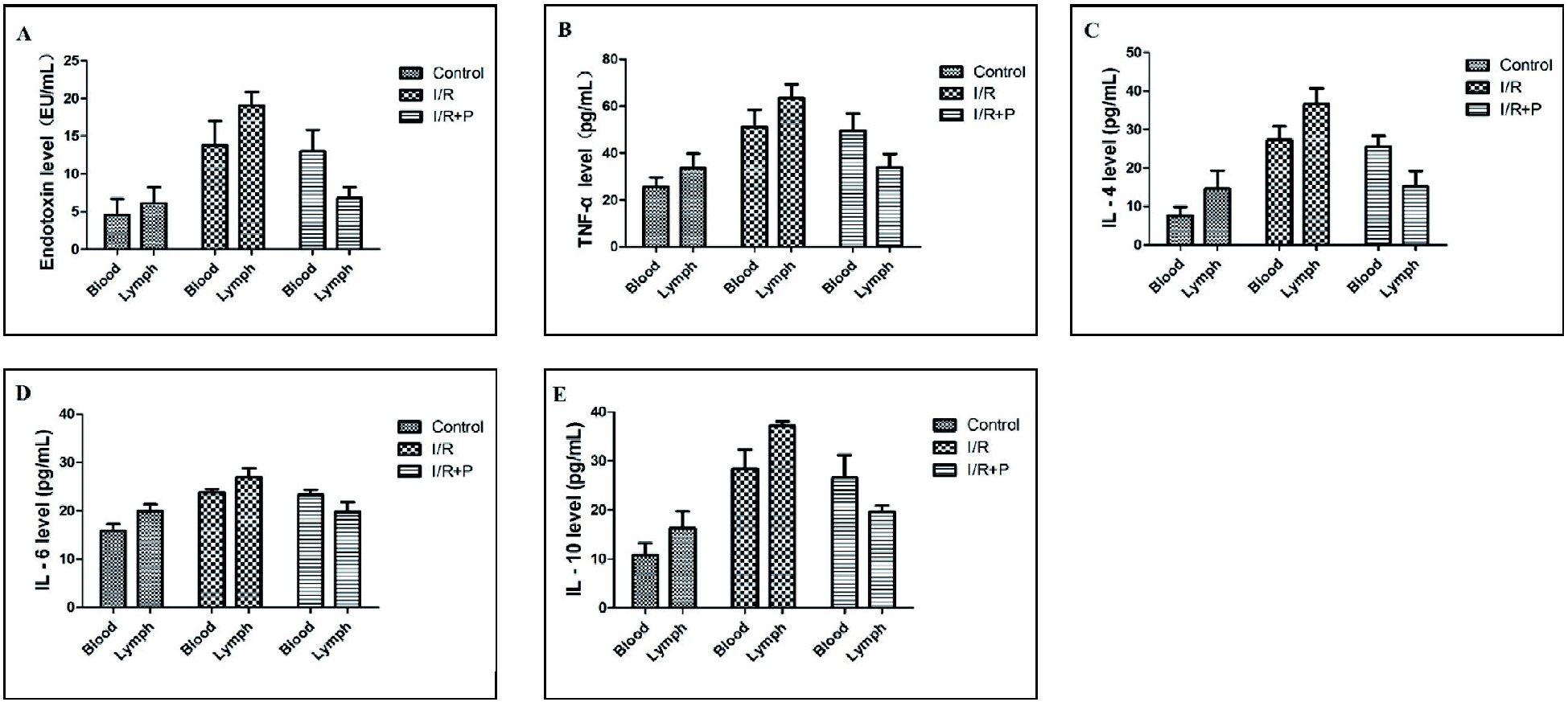
Levels of endotoxin (A), TNF-α (B), IL-4 (C), IL-6 (D), and IL-10 (E) in lymph fluid and plasma of the three groups (±s, n=8)

**Fig 2.**
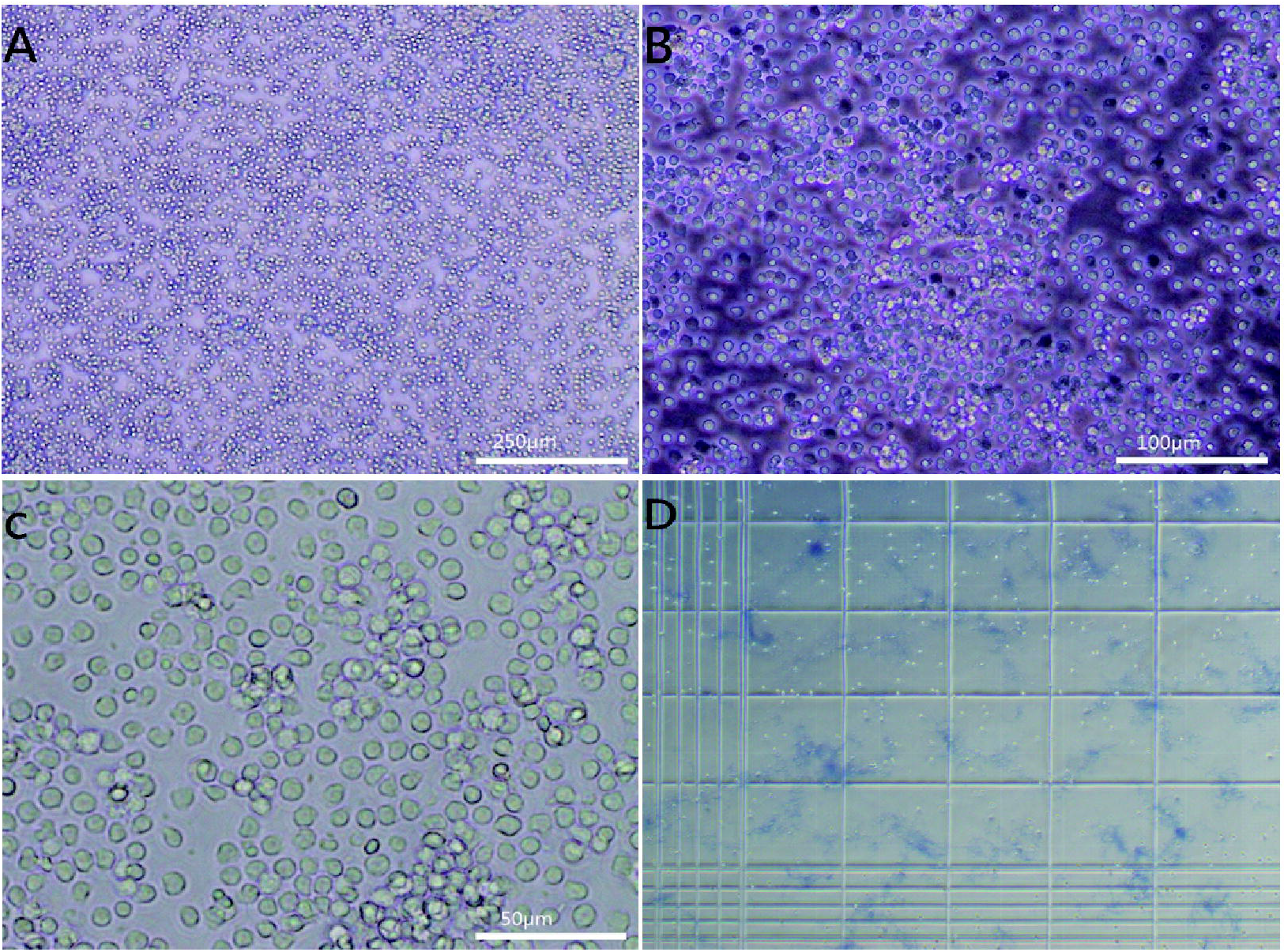
Monocytes were observed by inverted phase contrast microscope (A) magnification, ×40; (B) magnification, ×100; (C) magnification, ×200; (D) trypan blue rejection test

### Monocyte proliferation

Peripheral blood monocytes were co-cultured with lymph from each group to detect the monocyte proliferation rate. The proliferation rate of monocytes in the IRI group was significantly lower than that in the control group (*p* < 0.01). Compared with IRI group, the proliferation rate of monocytes in the IRI+GLP group was significantly increased (*p* < 0.01; Fig 3), showing that the lymph inhibited monocyte proliferation after IRI, and GLP promoted monocyte proliferation.

**Fig 3.**
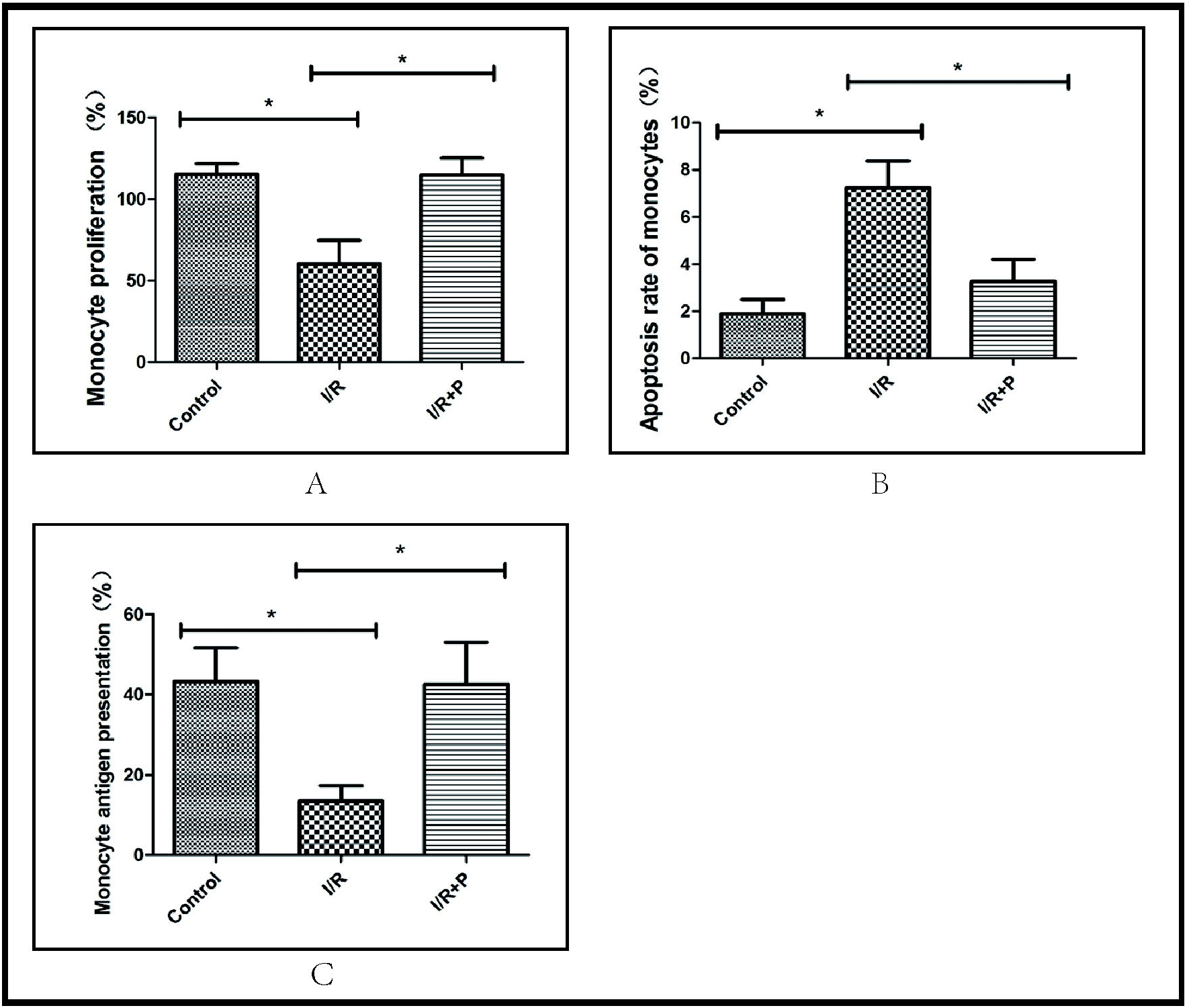
The proliferation (A), early apoptosis rate (B), and positive expression rate of MHC-II molecule (C) in monocytes in each group (±s, n=8) Note: * p<0.05

### Apoptotic rate of monocytes

Peripheral blood monocytes were co-cultured with lymph from each group to detect the early apoptotic rate of the monocytes. The apoptotic rate of monocytes in the IRI group was significantly higher than that in the control group (*p* < 0.01). Compared with IRI group, the apoptotic rate of monocytes in the IRI+GLP group was significantly decreased (*p* < 0.01; Figs 3 and 4). Thus, lymph promoted apoptosis of the monocytes after IRI, while GLP inhibited it.

**Fig 4.**
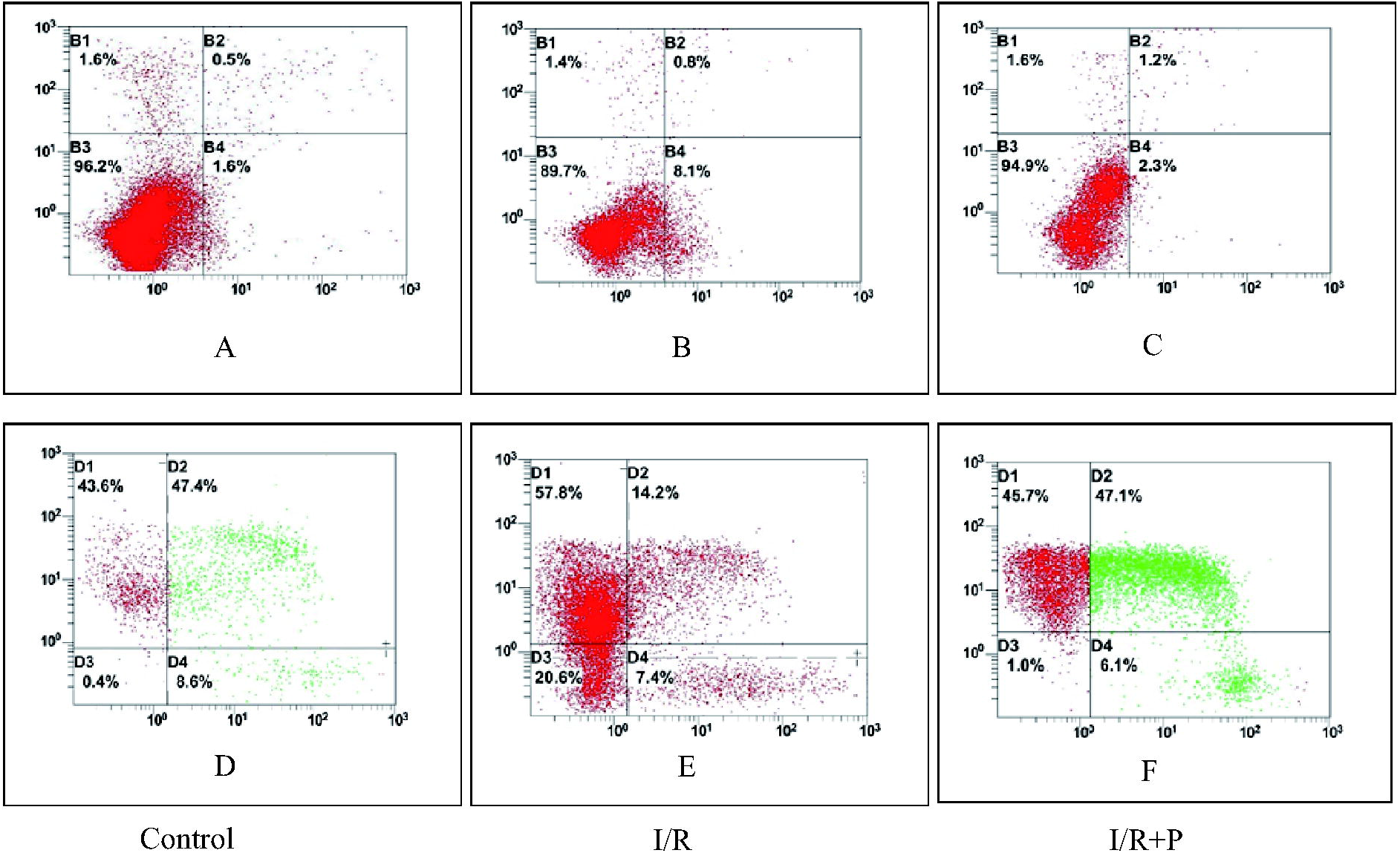
Apoptosis rate of monocytes and the positive expression rate of MHC-II molecule in monocytesin each group (n=8) Note: A, B, and C denote the apoptosis rate of monocytes: The horizontal staining is FITC-Annexin V; the vertical staining is PI. D, E, and F denote the positive expression rate of MHC-II molecule in monocytesin: OX-6 staining is used horizontally and OX-42 staining is used vertically.

### Monocyte antigen presentation

Peripheral blood monocytes were co-cultured with lymph from each group to detect the positive expression rate of MHC-II molecules in the monocytes. The positive expression rate of MHC-II molecules in the monocytes in the IRI group was significantly lower than that in the control group (*p* < 0.01). Compared with the IRI group, the positive expression rate of MHC-II molecules in the monocytes in the IRI+GLP group was significantly increased (*p* < 0.01; Figs 3 and 4), indicating that lymph inhibited the presentation of monocyte antigens after intestinal IRI, whereas GLP promoted it.

## Discussion

Researchers are paying increasing attention to the GL in the pathological process of MODS after gut IRI. Regulating the biological activity of the GL can reduce systemic inflammation and distant organ injury.[20] Rats are widely used as experimental animals to study the GL in lymph-related research.[21] In previous studies,[22] the rats were fasted for 6–12 hours pre-operation; however, the rats were not fasted in the current study. The lack of fasting did not affect our experimental process but instead made it smoother, possibly because more GLF was produced by enriching the gut-blood circulation, and more GLF was drained per unit time, which was more conducive to our experiments.

Inducing intestinal IRI in the experimental rodents increased the ITSs in the blood and lymph. Once intestinal IRI occurs, endotoxin, TNF-α and IL-6, which are important proinflammatory substances, promote inflammation. IL-4 and IL-10, which are critical anti-inflammatory substances, are increased in the blood and lymph to promote anti-inflammation when systemic inflammation is triggered. The molecular weights of these ITSs are endotoxin: 50–500 kDa, TNF-α: 17 kDa, IL-4: 18–19 kDa, IL-6: 21–30 kDa, and IL-10: 35–40 kDa. In blood purification, adsorption is the main method of scavenging small molecular substances with molecular weights >5 kDa. OXiris is an updated iteration product based on the structure of AN69 hydrogel, which can adsorb endotoxins and various proinflammatory and anti-inflammatory factors. In this experiment, endotoxin, TNF-α, IL-4, IL-6 and IL-10 levels in the plasma and lymph increased significantly after intestinal IRI in rats compared with those in the control group. OXiris purification significantly decreased the levels of these substances in the lymph. This indicated that GLP was effective and that GLP using oXiris can effectively remove endotoxins, TNF-a, IL-4, IL-6 and IL-10 and reduce their levels in the GLF. The plasma endotoxin, TNF-a, IL-6 and IL-10 levels did not significantly differ between the IRI and IRI+GLP groups after lymphatic reinfusion, possibly because the rats we used had less GL volume. Compared with the control group, lymphatic drainage volume in the IRI and IRI+GLP groups decreased significantly. Additionally, less GLF was transfused back in both groups postoperation. Although the levels of several ITSs transfused back in the lymph differed significantly between the two groups, less lymph was transfused back, resulting in no differences in the levels of several ITSs in the plasma of both groups after reinfusion. Therefore, using larger experimental animals (such as pigs or dogs) will yield more GLF for conducting additional experiments to compare the effects of GLP on several ITSs in the blood.

Toll-like receptor 4 (TLR-4) oligomerization is activated by endotoxins with the assistance of extracellular endotoxin-binding protein (LBP) and CD14 myeloid differentiation protein-2 (MD-2). TLR-4 transports stimulatory signals from lipopolysaccharides into cells, then further triggers cascade amplification of inflammation through two MyD88-dependent pathways to induce expression of proinflammatory factors, ultimately initiating the inflammatory response.[23] Inflammatory mediators (e.g., HGMB1), cytokines (e.g., TNF-α and IL-6) and leukotrienes (e.g., LTB4) play important roles in the GL theory of MODS, but the main mechanism is endotoxin activation in the monocyte-macrophage system. The endotoxin-activated monocyte-macrophage system mainly depends on TLR-4. TLR-4 transmits extracellular signals from endotoxins to cells via LBP, activates transcription factors such as nuclear factor κB (NF-κB) and activator protein-1 (AP-1) and further induces release of TNF-α, IL-6, interferon, and adhesion molecule (ICAM-1). An increase in inflammatory factors in the blood circulation increases the intestinal mucosa permeability, which makes endotoxins in the intestinal tract enter the interstitial space of the intestinal mucosa. These endotoxins are then absorbed into the systemic blood circulation and start a new round of monocyte-macrophage system activation, thus forming a cascade amplification waterfall effect on the inflammatory reaction.

MHC-II antigen-presenting molecules on the monocyte surface directly affect the intensity of the adaptive immune response. Because monocytes play a pivotal role in inflammation, we co-cultured peripheral blood monocytes with GLF to evaluate the effects of GLP on monocyte proliferation, apoptosis and antigen-presenting functions.

Researchers are paying increasing attention to the role of exosomes.[24] Kojima et al.[25] found exosomes in the GL of rats, which may come primarily from intestinal epithelial cells. Sakamoto et al.[26] reported that the lungs captured most of the exosomes injected into intestinal lymphatic ducts, suggesting that the lungs are the main target area to which the intestinal lymph carries exosomes. Kojima et al. further confirmed that exosomes significantly increased the expression of IL-8 homologues, which are the main cytokines for neutrophil aggregation, which promotes ALI development.[27] In addition, gut lymphatic exosomes in monocytes and macrophages can trigger NF-κB activation[25] and induce proinflammatory cytokine production[28] by monocytes and macrophages via the TLR-dependent signaling pathway, thus leading to lung injury. Exosomes mediate immunosuppression after injury and may provide a new direction for researching the intestinal lymphatic pathway.

## Conclusions

Using oXiris to perform GLP effectively removed ITSs from the GLF after IRI, thereby blocking the MODS process by regulating monocyte activity.

## Supporting information

Yes

Yes

Yes

## Declaration

### Ethics approval and consent to participate

The Ethics Committee of Experimental Animals and Use of Laboratory Animals of Zunyi Medical University approved this study.

### Consent to publish

All authors read and approved the manuscript version and agree with submitting to *Molecular Immunology* to be considered for publication. No ethical/legal conflicts are involved with the article.

### Availability of data and materials

The datasets used and/or analyzed during the current study available from the corresponding author on reasonable request.

### Competing interests

No ethical/legal conflicts are involved in this article.

### Funding

This study was supported by the Doctoral Research Initiation Fund of the Affiliated Hospital of Zunyi Medical College (2017-13), the Zunyi Medical College 2017 Academic New Seedling Cultivation and Innovative Exploration Fund (Qian Ke He Talents Platform [2017] 5733-019) and the Science and Technology Support Plan of Guizhou Province in 2019 (Qian Ke He Support [2019] 2834).

### Author contributions

Wei Zhang had full access to all data in the present study and accepts responsibility for data management and accuracy of the data analyses. Study concept and design: Wei Zhang, Jie Chen, and Meimei Shi. Acquisition and interpretation of data: Can Jin, Shucheng Zhang, and Juan Gu. Drafting of the manuscript: Can Jin, Shucheng Zhang, and Wei Zhang. Critical revision of the manuscript for important intellectual content: Meimei Shi and Wei Zhang. Administrative, technical, or material support: Wei Zhang and Shuncheng Zhang. Study supervision: Wei Zhang and Meimei Shi. All authors agree to submission of the final version of this manuscript. Wei Zhang is the study guarantor.

## Acknowledgements

The authors thank Professor Jidong Zhang and the members of his team. We thank Traci Raley, MS, ELS, from Liwen Bianji, Edanz Editing China (www.liwenbianji.cn/ac) for editing a draft of this manuscript.

## Additional Files

Additional Table 1. ITS levels in the GLF of rats in the three groups

Additional Table 2. ITS levels in the plasma of rats in the three groups

## References

1. Deitch EA, Xu D, Kaise VL, (2006) Role of the gut in the development of injury- and shock induced SIRS and MODS: the gut-lymph hypothesis, a review. Front Biosci 11: 520–528

2. Varela JE, Cohn SM, Diaz I, Giannotti GD, Proctor KG, (2003) Splanchnic perfusion during delayed, hypotensive, or aggressive fluid resuscitation from uncontrolled hemorrhage. Shock 20: 476–480

3. Moore EE, (1998) Mesenteric lymph: the critical bridge between dysfunctional gut and multiple organ failure. Shock 10: 415–416

4. Assimakopoulos SF, Triantos C, Thomopoulos K, Fligou F, Maroulis I, Marangos M, Gogos CA, (2018) Gut-origin sepsis in the critically ill patient: pathophysiology and treatment. Infection 46: 751–760

5. Watson AJ, Hughes KR, (2012) TNF-alpha-induced intestinal epithelial cell shedding: implications for intestinal barrier function. Annals of the New York Academy of Sciences 1258: 1–8

6. Moon HG, Cao Y, Yang J, Lee JH, Choi HS, Jin Y, (2015) Lung epithelial cell-derived extracellular vesicles activate macrophage-mediated inflammatory responses via ROCK1 pathway. Cell death & disease 6: e2016

7. Wohlauer MV, Moore EE, Harr J, Eun J, Fragoso M, Banerjee A, Silliman CC, (2011) Cross-transfusion of postshock mesenteric lymph provokes acute lung injury. The Journal of surgical research 170: 314–318

8. Zhang YM, Zhang SK, Cui NQ, (2014) Intravenous infusion of mesenteric lymph from severe intraperitoneal infection rats causes lung injury in healthy rats. World journal of gastroenterology 20: 4771–4777

9. Zhao ZG, Zhu HX, Zhang LM, Zhang YP, Niu CY, (2014) Mesenteric lymph drainage alleviates acute kidney injury induced by hemorrhagic shock without resuscitation. The Scientific World Journal 2014: 720836

10. Sambol JT, Lee MA, Caputo FJ, Kawai K, Badami C, Kawai T, Deitch EA, Yatani A, (2009) Mesenteric lymph duct ligation prevents trauma/hemorrhage shock-induced cardiac contractile dysfunction. Journal of applied physiology 106: 57–65

11. Sambol JT, White J, Horton JW, Deitch EA, (2002) Burn-induced impairment of cardiac contractile function is due to gut-derived factors transported in mesenteric lymph. Shock 18: 272–276

12. He GZ, Zhou KG, Zhang R, Wang YK, Chen XF, (2012) Impact of intestinal ischemia/reperfusion and lymph drainage on distant organs in rats. World journal of gastroenterology 18: 7271–7278

13. Zhang Y, Zhang S, Tsui N, (2015) Mesenteric lymph duct drainage attenuates acute lung injury in rats with severe intraperitoneal infection. Inflammation 38: 1239–1249

14. Tong H, Chen R, Yin H, Shi X, Lu J, Zhang M, Yu B, Wu M, Wen Q, Su L, (2016) Mesenteric Lymph Duct Ligation Alleviating Lung Injury in Heatstroke. Shock 46: 696–703

15. Zhang LM, Jiang LJ, Zhao ZG, Niu CY, (2014) Mesenteric lymph duct ligation after hemorrhagic shock enhances the ATP level and ATPase activity in rat kidneys. Renal failure 36: 593–597

16. de Jong PR, Gonzalez-Navajas JM, Jansen NJ, (2016) The digestive tract as the origin of systemic inflammation. Critical care 20: 279

17. Druml W, (2018) [Intestinal cross-talk: The gut as motor of multiple organ failure]. Medizinische Klinik, Intensivmedizin und Notfallmedizin 113: 470–477

18. Broman ME, Hansson F, Vincent JL, Bodelsson M, (2019) Endotoxin and cytokine reducing properties of the oXiris membrane in patients with septic shock: A randomized crossover double-blind study. PloS one 14: e0220444

19. Schwindenhammer V, Girardot T, Chaulier K, Gregoire A, Monard C, Huriaux L, Illinger J, Leray V, Uberti T, Crozon-Clauzel J, Rimmele T, (2019) oXiris(R) Use in Septic Shock: Experience of Two French Centres. Blood purification 47 Suppl 3: 1–7

20. Langness S, Costantini TW, Morishita K, Eliceiri BP, Coimbra R, (2016) Modulating the Biologic Activity of Mesenteric Lymph after Traumatic Shock Decreases Systemic Inflammation and End Organ Injury. PloS one 11: e0168322

21. Zhang HY, Besner GE, Feng JX, (2016) Antibody blockade of mucosal addressin cell adhesion molecule-1 attenuates proinflammatory activity of mesenteric lymph after hemorrhagic shock and resuscitation. Surgery 159: 1449–1460

22. Zhao Y, Zhang L, Han R, Si Y, Zhao Z, (2019) Intravenous injection of post-hemorrhagic shock mesenteric lymph induces multiple organ injury in rats. Experimental and therapeutic medicine 17: 1449–1455

23. Park JH, Jeong SY, Choi AJ, Kim SJ, (2015) Lipopolysaccharide directly stimulates Th17 differentiation in vitro modulating phosphorylation of RelB and NF-kappaB1. Immunology letters 165: 10–19

24. Raposo G, Stoorvogel W, (2013) Extracellular vesicles: exosomes, microvesicles, and friends. The Journal of cell biology 200: 373–383

25. Kojima M, Gimenes-Junior JA, Langness S, Morishita K, Lavoie-Gagne O, Eliceiri B, Costantini TW, Coimbra R, (2017) Exosomes, not protein or lipids, in mesenteric lymph activate inflammation: Unlocking the mystery of post-shock multiple organ failure. The journal of trauma and acute care surgery 82: 42–50

26. Sakamoto W, Masuno T, Yokota H, Takizawa T, (2017) Expression profiles and circulation dynamics of rat mesenteric lymph microRNAs. Molecular medicine reports 15: 1989–1996

27. Hu R, Chen ZF, Yan J, Li QF, Huang Y, Xu H, Zhang XP, Jiang H, (2015) Endoplasmic Reticulum Stress of Neutrophils Is Required for Ischemia/Reperfusion-Induced Acute Lung Injury. Journal of immunology 195: 4802–4809

28. Bonjoch L, Casas V, Carrascal M, Closa D, (2016) Involvement of exosomes in lung inflammation associated with experimental acute pancreatitis. The Journal of pathology 240: 235–245

