## Supplementary material for "Gut lymph purification regulates monocyte activity in rats with ischemia-reperfusion injury-induced sepsis": Yes

**Table 1. The comparison between IRI, IRI combined GLP and control group for lymphatic drainage (**
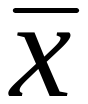
**±s, n=8)**

| **Group** | **Drainage（mL）** | **Retransmission（mL）** |
| --- | --- | --- |
| **Control** | 1.79±0.542 | 1.075±0.292 |
| **I/R** | 1.488±0.155^*^ | 0.763±0.177^*^ |
| **I/R+P** | 1.475±0.191^*^ | 0.575±0.175^*^ |

Note:^*^Compared with the control group，*p*＜0.05
