## Supplementary material for "Gut lymph purification regulates monocyte activity in rats with ischemia-reperfusion injury-induced sepsis": Yes

Table 2. Levels of ITSs in the GLF of rats in the three groups (
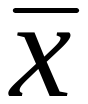
±s, n=8)

| Group | Endotoxin（EU/ml） | TNF-α（pg/mL） | IL-4（pg/mL） | IL-6（pg/mL） | IL-10（pg/mL） |
| --- | --- | --- | --- | --- | --- |
| Control | 6.109±2.093 | 33.542±6.295 | 14.597±4.725 | 19.923±1.444 | 16.242±3.518 |
| I/R | 19.027±1.806* | 63.5±5.757* | 36.575±4.071* | 26.964±1.804* | 37.190±0.843* |
| I/R+P | 6.779±1.45.6# | 33.952±5.668 # | 15.174±4.083# | 19.750±1.990# | 19.573±1.290# |

Note:^*^Compared with the control group，*p*＜0.05；^#^Compared with the I/R group，*p*＜0.05
