## Supplementary material for "Gut lymph purification regulates monocyte activity in rats with ischemia-reperfusion injury-induced sepsis": Yes

Table 3. Levels of ITSs in plasma of rats in the three groups (
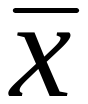
±s, n=8)

| Group | Endotoxin（EU/ml） | TNF-α（pg/mL） | IL-4（pg/mL） | IL-6（pg/mL） | IL-10（pg/mL） |
| --- | --- | --- | --- | --- | --- |
| Control | 4.551±2.150 | 25.595±4.019 | 7.603±2.233 | 15.809±1.453 | 10.800±2.429 |
| I/R | 13.799±3.17 * | 51.153±7.290* | 27.270±3.546* | 23.790±0.691* | 28.379±3.911* |
| I/R+P | 13.020±2.781* | 49.422±7.538* | 25.540±2.835*# | 23.300±1.042* | 26.587±4.633* |

Note:^*^Compared with the control group，*p*＜0.05；^#^Compared with the I/R group，*p*＜0.05
