## Supplementary material for "Gut lymph purification regulates monocyte activity in rats with ischemia-reperfusion injury-induced sepsis": No

### Please wait...

If this message is not eventually replaced by the proper contents of the document, your PDF viewer may not be able to display this type of document.

You can upgrade to the latest version of Adobe Reader for Windows®, Mac, or Linux® by visiting  http://www.adobe.com/products/acrobat/readstep2.html.

For more assistance with Adobe Reader visit  http://www.adobe.com/support/products/acrreader.html.

Windows is either a registered trademark or a trademark of Microsoft Corporation in the United States and/or other countries. Mac is a trademark of Apple Inc., registered in the United States and other countries. Linux is the registered trademark of Linus Torvalds in the U.S. and other countries.
